# Spatial cell type mapping of the oligodendrocyte lineage in the mouse juvenile and adult CNS with in situ sequencing

**DOI:** 10.1101/2021.06.04.447052

**Authors:** Markus M. Hilscher, Christoffer Mattsson Langseth, Petra Kukanja, Chika Yokota, Mats Nilsson, Gonçalo Castelo-Branco

## Abstract

Oligodendrocytes show transcriptional heterogeneity but the regional and functional implications of this heterogeneity are less clear. Here, we apply *in situ* sequencing (ISS) to simultaneously probe the expression of 124 marker genes of distinct oligodendrocyte populations, providing comprehensive maps of corpus callosum, cingulate, motor and somatosensory cortex in the brain, as well as gray (GM) and white matter (WM) regions in the spinal cord, at juvenile and adult stages. We systematically compare abundances of these populations and investigate the neighboring preference of distinct oligodendrocyte populations. As previously described, we observed that oligodendrocyte lineage progression is more advanced in the juvenile spinal cord compared to the brain. Additionally, myelination is ongoing in the adult corpus callosum while it is mostly completed in the cortex. Interestingly, we found a medial-to-lateral gradient of oligodendrocyte lineage progression in the juvenile cortex, which could be linked to arealization, as well as a deep-to-superficial gradient with mature oligodendrocytes preferentially accumulating in the deeper layers of the cortex. We observed differences in abundances and population dynamics over time between GM and WM regions in the brain and spinal cord, indicating regional differences within GM and WM. We also found that oligodendroglia populations’ neighboring preferences are altered from the juvenile to the adult CNS. Thus, our ISS dataset reveals spatial heterogeneity of the oligodendrocyte lineage progression in the brain and spinal cord, which could be relevant to further investigate functional heterogeneity of oligodendroglia.

## INTRODUCTION

Classically, oligodendrocytes (OLs) have been considered a homogeneous population in the central nervous system (CNS). Recent advances in single cell RNA sequencing (sc-RNA seq) have allowed systematic studies of the transcriptome in individual cells, which changed the paradigm of cellular classification (Yuste et al, 2020). At least twelve distinct OL populations can be distinguished in the CNS (Marques et al, 2016). If these transcriptionally distinct populations are indeed different cell types or rather states remains a recurring question in the field (Foerster et al, 2019) and their spatial distribution could give further insights into the heterogeneity of OLs.

The OL lineage presents itself as a continuum of OPCs (oligodendrocyte precursor cells), COPs (committed OPCs), NFOLs (newly formed OLs), MFOLs (myelin-forming OLs) and MOLs (mature OLs) (Marques et al, 2016). In our previous study, using RNASCOPE *in situ* hybridization and a limited number of markers, we have shown that OPC, MOL2 and MOL5/6 take spatial preference in the CNS and the abundance changes with age (Floriddia et al, 2020). More precisely in the neocortex and corpus callosum, the OPC population decreases with age while the MOL5/6 population increases in both regions (Floriddia et al, 2020). Also in the spinal cord the MOL5/6 population increases with age and it shows spatial preference for gray matter, while the MOL2 population shows spatial preference for white matter (Floriddia et al, 2020).

Most recently, novel spatial transcriptomics methods have been developed that can map the location of each gene and each cell (Strell et al, 2019; Asp et al, 2020). Multiple *ex situ* and *in situ* spatial transcriptomics technologies are now available. *In situ* sequencing (ISS) is a targeted approach that allows the precise mapping of genes in its natural context (Ke et al, 2013). Here, we applied pciSeq (probabilistic cell typing by *in situ* sequencing; (Qian et al, 2019)) to target OLs for the first time, leveraging sc-RNA seq data to assign detected transcripts to segmented cells and subsequently, cells to cell types. We used the full advantages of ISS by probing 124 marker genes that characterize all twelve previously described populations of the OL lineage and the VLMCs (vascular and leptomeningeal cells) (Marques et al, 2016). We investigated multiple brain and spinal cord tissue sections and found region- and age-related differences of the oligodendrocyte lineage progression, uncovering differences in timing of oligodendrocyte differentiation and myelination in these areas.

## RESULTS

### *In Situ* sequencing allows detection of major OL populations

We performed *in situ* sequencing (ISS) in juvenile (P20) and adult (P60) mouse brain sections as well as thoracic spinal cord sections (6+6 hemisected coronal brain including neocortex (CTX) and corpus callosum (CC) and 4+4 spinal cord sections, from 8 mice in total) and we quantified the OL lineage populations in their white matter (WM) vs. gray matter (GM), respectively. The gene maps of CC (WM) in a P60 brain section are shown in **Figure 1A**, with the colored symbols representing unique transcripts. Using pciSeq (Qian et al, 2019), we assigned detected transcripts to segmented cells and subsequently, based on their genetic signature cells to cell populations. Thereby, each cell is represented as a pie chart, with the angle of each slice being proportional to the cell type probability. The highest probability in the pie chart defines the final cell population. The mean probability for the OL assignments in P20 and P60 brains was 77%. Only cells that express *Plp1* as a PAN marker for OLs were considered for downstream analysis. Example cells that are cells with the median count of genes per population are shown in **Figure 1B**. On average, 14.6 ± 2.1 transcripts and 10.0 ± 0.1 distinctive genes per cell were measured in P20 brains and 17.4 ± 2.7 transcripts and 9.8 ± 0.1 distinctive genes per cell were measured in the OL populations in P60 brains (**Supplementary Figure 1A, B**). Heatmaps of the log2 gene expression per cell population are shown in **Supplementary Figure 1C**, emphasizing that many OL-associated genes are shared between OL populations but that the composition of genes per OL population is unique, allowing us to map these populations with pciSeq.

**Figure 1:**
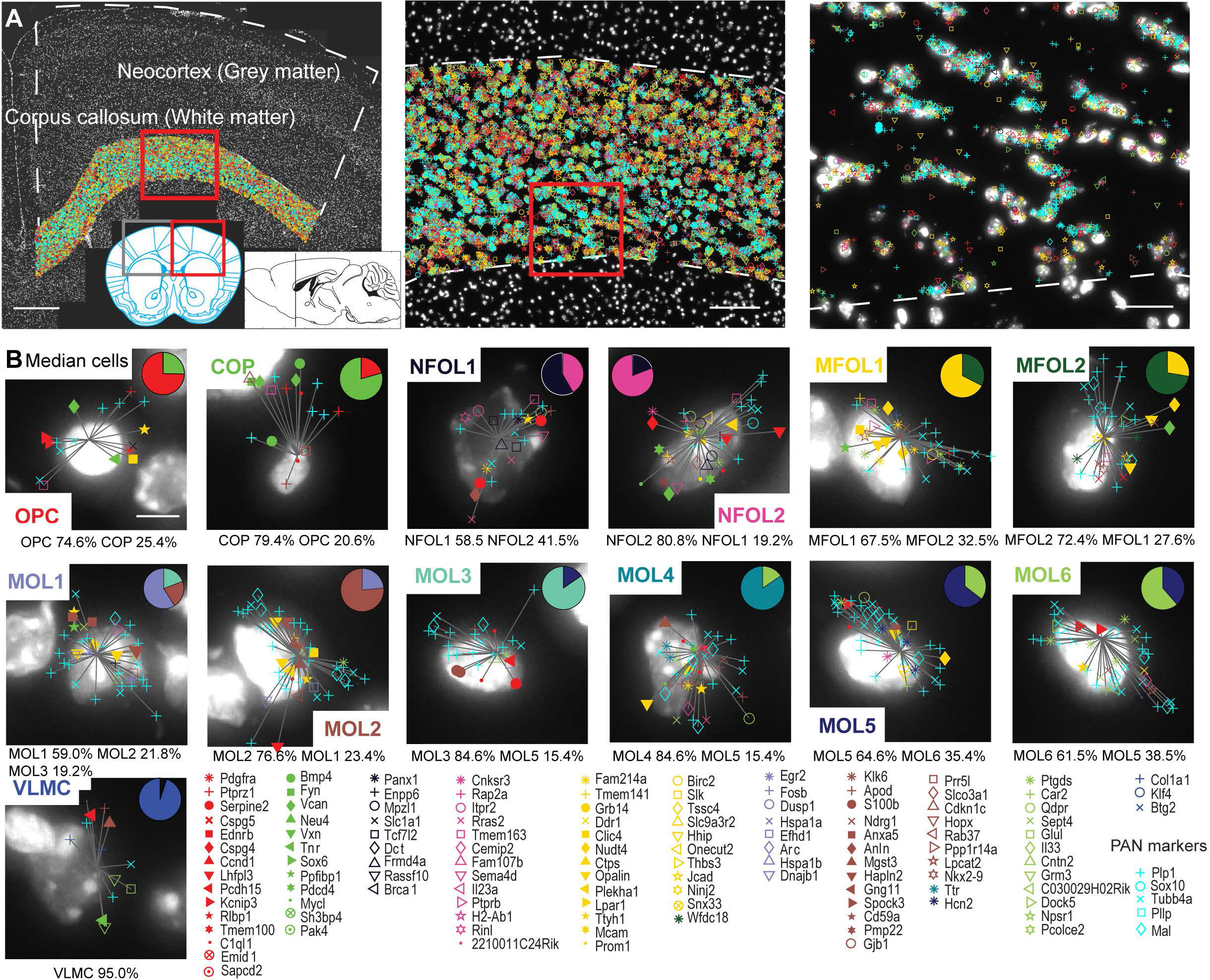
Oligodendrocyte types in neocortex (CTX), grey matter (GM) and corpus callosum (CC), white matter (WM) **a**, Maps of 124 genes in coronal mouse brain sections (P60) targeted by *In Situ* sequencing, here highlighting the CC. Three different zoom levels are shown with white dashed lines denoting the outlines for CTX and CC. The red boxes mark the positions of the zoom-ins. The scale bars are 500 μm (left), 100 μm (middle) and 15 μm (right). **b**, Individual example cells of P60 CC and their gene assignments to cells are shown. The 124 gene symbols and gene names are shown below the images of the cells. The cells are classified based on single cell RNA-sequencing (scRNA-seq) data and represented as pie charts where the angle of each slice is proportional to the likelihood of the cell being of a certain type and the size of the pie chart is indicative of the number of transcripts. Examples shown are median cells for each cell population, i.e. cells with the median count of the number of transcripts being assigned within each type. The probability for each cell type is listed below the example cells. The scale bar is 5 μm.

Representative cell maps of P20 and P60 OL populations in CTX (GM) and CC (WM) are shown in **Figure 2A, B**. The size of each pie chart is indicative of the number of assigned transcripts. We also generated mean pie charts counting how often a certain OL population was called with other OL populations (**Figure 2D, E**). The mean pie charts per OL populations over the brain tissues (n=6 tissue sections per postnatal age) reflect the progression of OL lineage very well, with OPCs often being shared with COPs, NFOLs with MFOLs and MFOLs with MOLs. The relatedness of the six MOL populations can also be seen in the mean confusion matrix of the cell populations during the probabilistic assignment (**Supplementary Figure 2**).

**Figure 2:**
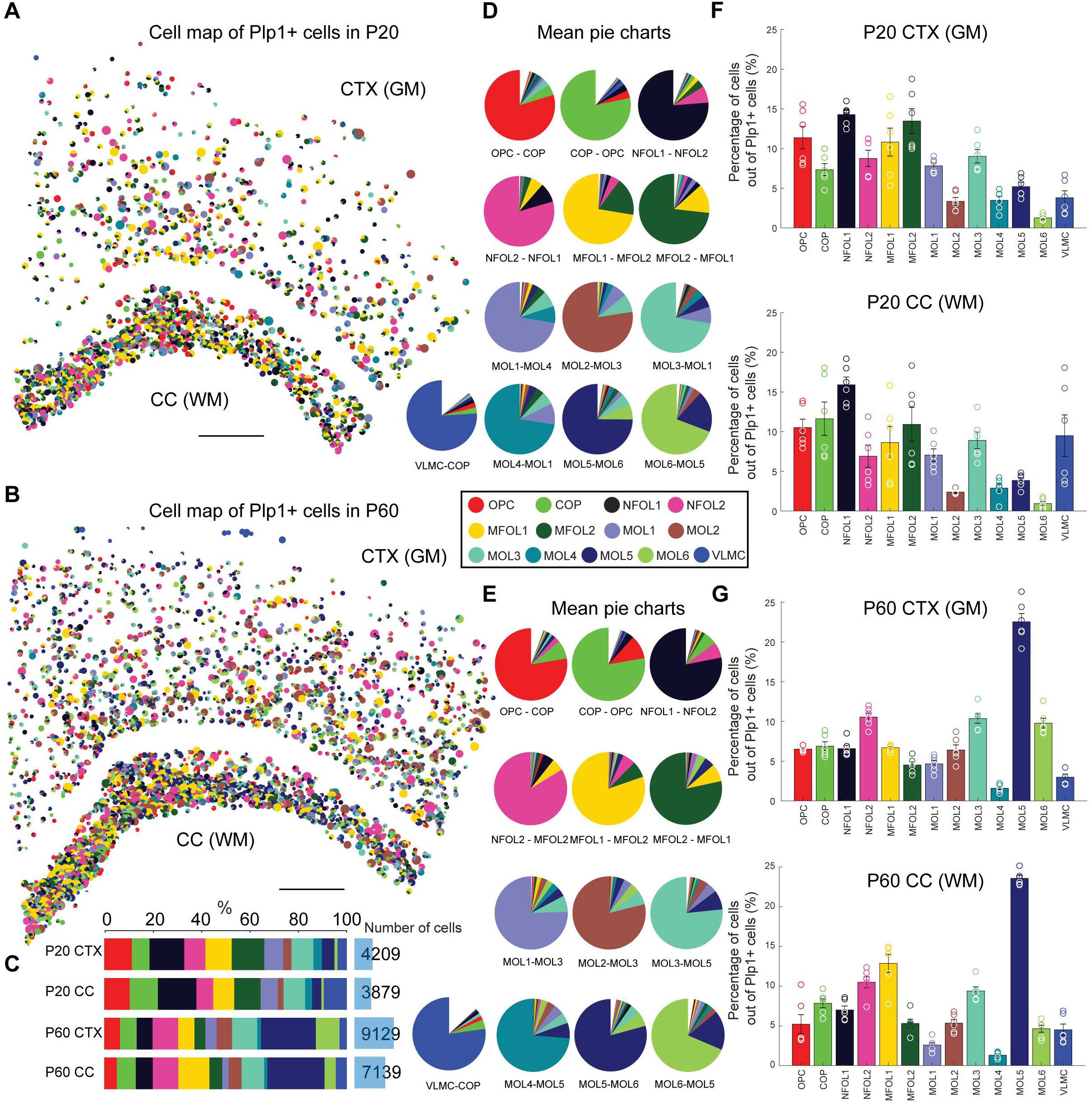
Oligodendrocyte maps and abundances in P20 and P60 CTX (GM) and CC (WM) **a**, Spatial map of Plp1+ cell types in a P20 coronal mouse brain section including neocortex (CTX) and corpus callosum (CC). The scale bar is 500 μm. **b**, Same as **a** for P60. **c**, The number of oligodendrocytes as well as the relative distribution of the oligodendrocyte types per age condition and brain region. **d**, Mean pie charts for P20 (CTX and CC joined), generated by counting how often a certain OL population was called with other OL populations. Note: Considering the top pies within the pie charts (caption below), the charts highlight the oligodendrocyte lineage from OPC to MOL6. **e**, Same as **d** for P60. **f**, Mean relative number of OL types for P20 CTX (top) and CC (bottom). The counts are normalized as counts per class divided by the total counts of Plp1+ cells per tissue for CTX and CC, respectively. Circles represent the counts in individual sections, error bars represent the standard error of the mean. We mapped on average 76 ± 8 OPC, 50 ± 7 COP, 98 ± 10 NFOL1, 65 ± 15 NFOL2, 82 ± 21 MFOL1, 100 ± 23 MFOL2, 55 ± 8 MOL1, 25 ± 7 MOL2, 60 ± 5 MOL3, 24 ± 4 MOL4, 35 ± 3 MOL5, 8 ± 1 MOL6 and 24 ± 3 VLMC in CTX and 64 ± 8 OPC, 67 ± 3 COP, 107 ± 23 NFOL1, 52 ± 16 NFOL2, 67 ± 23 MFOL1, 82 ± 26 MFOL2, 43 ± 6 MOL1, 15 ± 2 MOL2, 52 ± 5 MOL3, 21 ± 6 MOL4, 27 ± 6 MOL5, 7 ± 2 MOL6 and 47 ± 6 VLMC in CC (n=6 hemisected tissue sections from 3 mice). **g**, Same as **f** for P60. We mapped on average 99 ± 9 OPC, 102 ± 7 COP, 99 ± 10 NFOL1, 160 ± 16 NFOL2, 103 ± 11 MFOL1, 68 ± 9 MFOL2, 71 ± 9 MOL1, 100 ± 15 MOL2, 158 ± 18 MOL3, 25 ± 4 MOL4, 342 ± 37 MOL5, 152 ± 21 MOL6 and 43 ± 5 VLMC in CTX and 62 ± 18 OPC, 93 ± 7 COP, 84 ± 8 NFOL1, 125 ± 10 NFOL2, 153 ± 14 MFOL1, 63 ± 7 MFOL2, 31 ± 4 MOL1, 64 ± 6 MOL2, 111 ± 4 MOL3, 15 ± 2 MOL4, 280 ± 8 MOL5, 55 ± 5 MOL6 and 54 ± 9 VLMC in CC (n=6 hemisected tissue sections from 3 mice).

### Myelination advances earlier in GM (CTX) than in WM (CC) in the brain

In total, we captured 4209 OLs in P20 CTX, 3879 OLs in P20 CC, 9129 OLs in P60 CTX and 7139 OLs in P60 CC, (n=6 tissue sections per postnatal age, **Figure 2C**). For P20, we captured a tissue area of 3.6 ± 0.3 mm^2^ of CTX and a tissue area of 0.9 ± 0.1 mm^2^ of CC, yielding an OL density of 193.5 ± 31.8 cells/mm^2^ in CTX and an OL density of 750.4 ± 82.8 cells/mm^2^ in CC (p = 0.0003, n=6 tissue sections). For P60, we captured a tissue area of 8.9 ± 0.6 mm^2^ of CTX and a tissue area of 2.0 ± 0.1 mm^2^ of CC, with an OL density of 170.1 ± 48.2 cells/mm^2^ in CTX and an increased OL density of 608.6 ± 190.4 cells/mm^2^ in CC (p = 0.05, n=6 tissue sections), as expected for gray and white matter regions. As expected, we also observed a 4-5x higher density of the different OL populations in CC compared to CTX (**Supplementary Figure 3**). Counting the relative cell occurrences (number of cells per Plp1+ cells) in the P20 brain tissue sections, when myelination is actively ongoing, shows that OPC, COP, NFOL1 and MFOL2 are most abundant in CTX and CC (**Figure 2F**), while at P60 MOL5 are most abundant in CTX and CC and MOL1 and MOL4 populations are a minority (**Figure 2G**).

Next, we compared the abundances of OL populations at P20 and P60, between GM (CTX) and WM (CC) by calculating how much the OL population in the prevalent region is increased compared to the non-prevalent region. In P20, we found that NFOL2 (25 ± 8% increased in GM versus WM), MFOL1 (25 ± 7%) and MFOL2 (22 ± 8%) are more abundant in GM (CTX) compared to WM (CC), while there is an enrichment of COPs (31 ± 8%) and VLMC (56 ± 4%) in CC compared to CTX (p = 0.03, p = 0.004, p = 0.02, p = 0.05, p = 0.02, respectively; n=6 tissue sections) (**Figure 3A, left**). Interestingly, OPCs were not differentially distributed between these two regions at P20.

**Figure 3:**
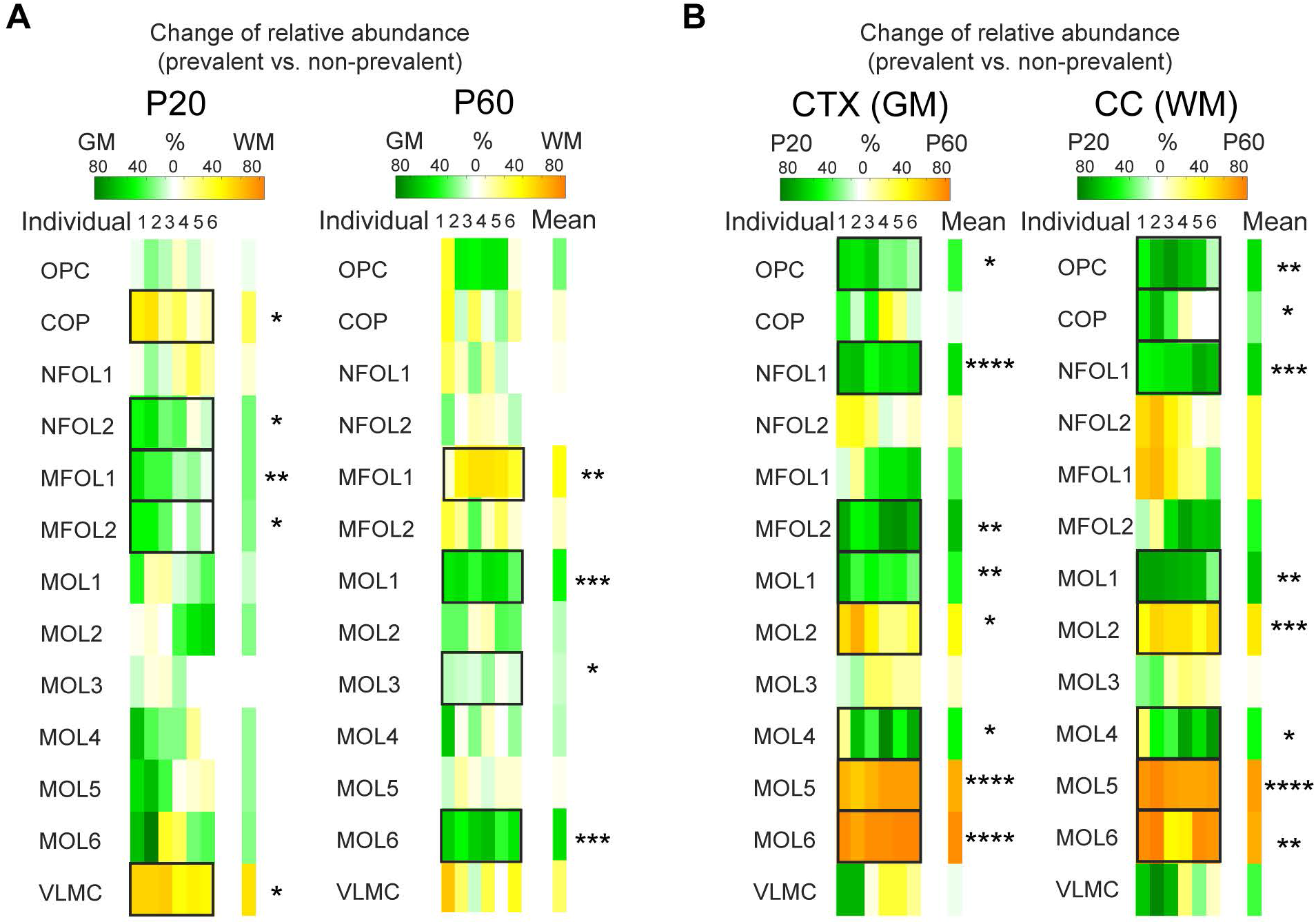
Change of relative oligodendrocyte abundances in P20 and P60 CTX (GM) and CC (WM) **a**, Change of relative abundances of oligodendrocytes between CTX (GM) and CC (WM) for P20 (left) and P60 (right). Changes were calculated on how much the OL population in the prevalent region is increased compared to the non-prevalent region. Statistical comparisons were determined using ANOVA and post-hoc test with Tukey correction (* p < 0.05, ** p < 0.01, *** p < 0.001, and **** p < 0.0001). **b**, Change of relative abundances of oligodendrocytes between P20 and P60 for CTX (left) and CC (right).

At P60, comparing the relative occurrences of OLs over all six tissue sections, we found MFOL1 to be more abundant in WM (CC) compared to GM (CTX) (45 ± 7%), whereas mature OL populations such as MOL1 (45 ± 4%), MOL3 (9 ± 3%) and MOL6 (52 ± 4%) were more enriched in CTX (p = 0.004, p = 0.0004, p = 0.02, p = 0.0002, respectively; n=6 tissue sections) (**Figure 3A, right**). These data suggest that myelination might be still ongoing in the CC (WM) at P60, while being mostly completed in the CTX (GM), which is more populated with mature OLs.

### Decrease of MOL1 and increase of MOL2, MOL5 and MOL6 with age in both GM (CTX) and WM (CC) in the brain

Comparing the distributions of OL populations between the juvenile and adult brain shows a relative decrease of OPCs (38 ± 8%), NFOL1 (54 ± 3%) and MFOL2 (64 ± 6%) with age in the CTX (**Figure 3B, left**). Interestingly, mature populations as MOL1 (40 ± 6%) and MOL4 (44 ± 15%) were also more abundant at P20 CTX compared to P60 CTX (p = 0.02, p ≤ 0.0001, p = 0.003, p = 0.002, p = 0.03). MOL1 expresses Egr2/Krox20, which was previously identified as a transcription factor required for the early myelinating phase of Schwann cells (Topilko et al, 1994). Thus, MOL1 might be an early myelinating population that later transitions into other mature populations. MOL1 is also characterized by the expression of genes involved in response to stress (Marques et al, 2016; Floriddia et al, 2020) and this reduction might reflect targeted sensitivity or even apoptosis of this OL population. In contrast MOL2 (45 ± 9%), MOL5 (77 ± 3%) and MOL6 (87 ± 2%) in CTX were more abundant with age (p = 0.01, p ≤ 0.0001, p ≤ 0.0001, respectively; n=6 tissue sections per age, **Figure 3B, left**).

We observed very similar trends in the brain WM (CC), where there was a decrease with age of OPC (50 ± 10%), COP (22 ± 12%), NFOL1 (55 ± 4%), MOL1 (61 ± 8%) and MOL4 (41 ± 16%) and an increase of MOL2 (54 ± 3%), MOL5 (84 ± 1%) and MOL6 (76 ± 8%) (p = 0.008, p = 0.05, p = 0.0005, p = 0.004, p = 0.03, p = 0.0006, p ≤ 0.0001, p = 0.002, respectively; n=6 tissue sections per age, **Figure 3B, right**). Thus, our data suggests that the terminal differentiation into specific mature MOLs occurs at a similar dynamics in GM (CTX) and WM (CC) in the brain (**Figure 3B**), despite the different pace (**Figure 3A**).

### Enrichment of specific OL populations at distinct areas and layers in the CTX

Specific populations of neurons and even fine transcriptomic cell types are differentially distributed in six distinct layers and areas in the CTX (Zeisel et al, 2015; Tasic et al, 2018). We thus investigated whether the same applies to OLs. First, we plotted the kernel density estimates for each of the twelve OL populations, highlighting the spatial preference of certain OL populations (**Figure 4A**). We further segmented the sections according to Allen Brain Atlas into primary somatosensory cortex (S1), primary motor cortex (M1), secondary motor cortex (M2), cingulate cortex area1 (Cg1) and cingulate cortex area2 (Cg2) (Lein et al, 2007). At P20, we found NFOL1, MFOL1 and MFOL2 being more abundant in the medial regions M2, Cg1 and Cg2, while OPC and MOL1 were decreased in Cg2 (OPC: p = 0.004, NFOL1: p = 0.003, p = 0.002, MFOL1: p = 0.005, p = 0.005, MFOL2: p = 0.005, p = 0.008, MOL1: p = 0.002, MOL5: p = 0.002, p = 0.003, respectively; n=6 tissue sections per condition, **Figure 4B, top**). Mature OL populations were equally distributed in the different areas, with the exception of MOL5, which was more prevalent in S1 than in other regions and less prevalent in M2 than in other regions (p = 0.002, p = 0.003, respectively). Thus, our ISS analysis indicates that the different areas in the CTX are in different stages of OL lineage progression in the juvenile brain, with areas such as somatosensory cortex being more advanced in comparison to motor cortex and cingulate cortex.

**Figure 4:**
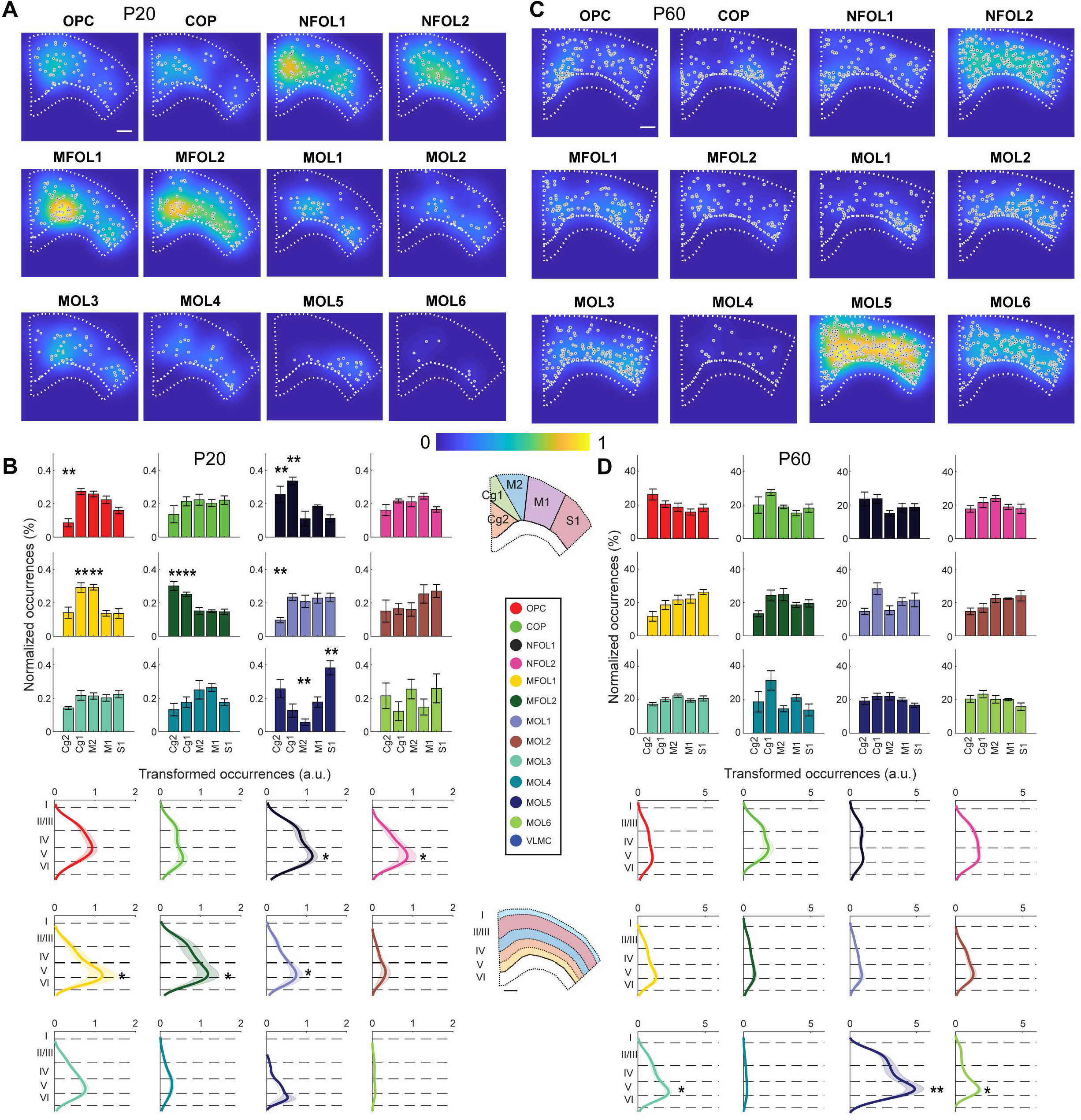
Density of oligodendrocyte types within P20 and P60 CTX (GM) **a**, Heatmaps of density of cells for each oligodendrocyte type, normalized by the maximum density over all types. The scale bar is 500 μm. **b**, Top: Mean normalized occurrences of oligodendrocyte populations in primary somatosensory cortex (S1), primary motor cortex (M1), secondary motor cortex (M2), cingulate cortex area1 (Cg1) and cingulate cortex area2 (Cg2). Bottom: Mean cortical depth profiles with the transparent shades representing the standard error of the mean. Dashed lines denote layers I, II/III, IV, V and VI. X-axis (top) and Y-axis (bottom), respectively, represent occurrences after smoothening non-binned data with a kernel. Statistical comparisons were determined using ANOVA and post-hoc test with Tukey correction (* p < 0.05, ** p < 0.01, *** p < 0.001, and **** p < 0.0001). **c**, Same as a for P60 CTX. **d**, Same as b for P60 CTX.

We further segmented the five neocortical layers and found that there are less OLs in layer I compared to deeper layers (I vs. II/III: p = 0.001; I vs. IV: p = 0.0008; I vs. V: p = 0.0006; I vs. VI: p = 0.0003, n=6 tissue sections per condition), in accordance with the literature (McGee et al, 2005; Vincze et al, 2008). In addition, there were less OLs in layer II/III and layer IV compared to layer V and layer VI, respectively (II/III vs. V: p = 0.001; II/III vs. VI: p = 0.004; IV vs. V: p = 0.03; IV vs. VI: p = 0.05, n=6 tissue sections per condition). OLs associated with early stages of differentiation and myelination, such as NFOL1, NFOL2, MFOL1 and MFOL2 in P20 brains exposed an increased occurrence in the infragranular layers, peaking at layer V and VI (p = 0.01, p = 0.01, p = 0.02, p = 0.03, p = 0.03, respectively; n=6 tissue sections per condition, **Figure 4B, bottom**). These data could suggest that the cellular environment at layer V and VI might actively be involved with the induction of OL differentiation and myelination. Interestingly, MOL1 was also enriched in this layer at P20 (p = 0.03, respectively; n=6 tissue sections, **Figure 4B, bottom**). OL differentiation and OL stress/apoptosis are closely connected (Sun et al, 2018), and our data suggests that the local environment at the cortical layer V at P20 might be amenable for both these processes.

We performed the same analysis on the P60 brain sections (**Figure 4C**). In contrast to P20, we did not observe any enrichment in the different cortical areas at P60 (**Figure 4D, top**), suggesting that the process of differentiation, myelination and subtype specification was completed at this stage (although it is possible that specified MOL can still form new myelin sheaths later in late adulthood). Similar to P20, there were less OLs in layer I compared to deeper layers (I vs. II/III: p = 0.02; I vs. IV: p = 0.02; I vs. V: p = 0.001; I vs. VI: p = 0.001, n=6 tissue sections per condition) as well as less OLs in layer II/III and layer IV compared to layer V and layer VI, respectively (II/III vs. V: p = 0.03; II/III vs. VI: p = 0.02; IV vs. V: p = 0.05; IV vs. VI: p = 0.05, n=6 tissue sections per condition).

In P60 cortical brain sections the early differentiating/myelinating OLs and MOL1 presented a more uniform distribution when compared to P20. Moreover, there was an increased abundance in deeper regions, particularly again at layer V/VI, of mature OL populations such as MOL3, MOL5 and MOL6 populations (p = 0.05, p = 0.008, p = 0.04, respectively; n=6 tissue sections, **Figure 4D, bottom**). Thus, our data suggest that layer V and VI are particularly enriched for myelinating OL at P20, especially in medial regions, while at P60 these layers are enriched in specific mature OL populations.

### Differential dynamics and abundances in brain and spinal cord WM and GM

Next, we investigated the occurrence of the OL populations in the P20 and P60 spinal cord. The gene map of a thoracic P60 spinal cord section is shown in **Figure 5A, left**, with colored symbols representing the unique transcripts and the segmented gray matter (GM) and white matter (WM) cell maps are shown in **Figure 5A, right**. Similar to the brain we also assessed cell density by heatmaps (**Figure 5B**). Note that we splitted the heatmaps for GM and WM to allow comparisons and highlight the cellular densities in the dorsal funiculus region of WM.

**Figure 5:**
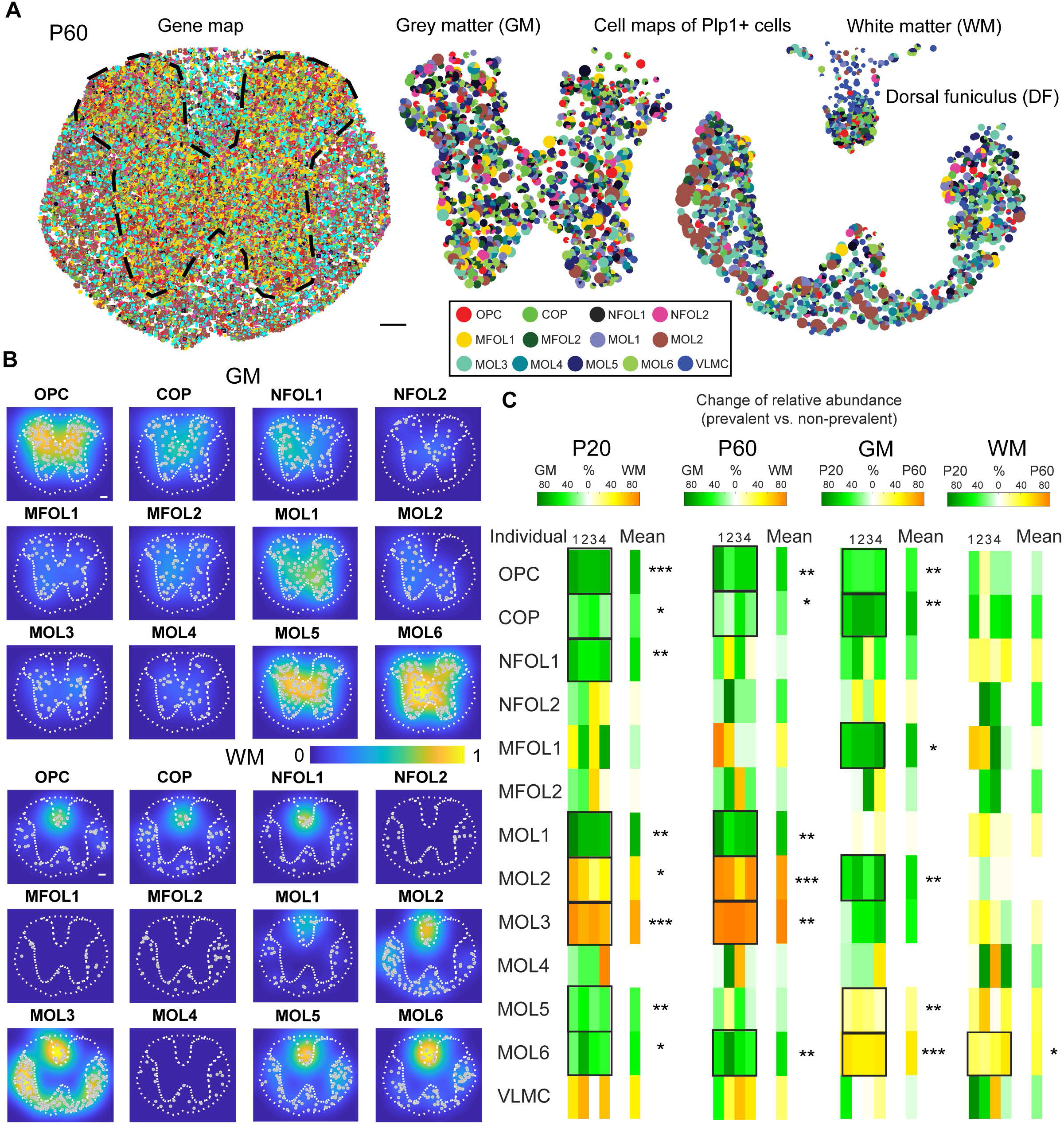
Oligodendrocyte types in spinal cord gray matter (GM) and white matter (WM) **a**, Maps of 124 genes (left) and 13 cell types (middle and right) in spinal cord sections (P60) targeted by In Situ sequencing. The dashed line in the gene map highlights the separation between gray matter (GM) and white matter (WM). The scale bar is 100 μm. **b**, Heatmaps of density of cells for each oligodendrocyte type, normalized by the maximum density over all types. The scale bar is 100 μm. **c**, Change of relative abundances of oligodendrocytes between GM and WM for P20 and P60 (left) as well as change of relative abundances of oligodendrocytes between P20 and P60 for GM and WM (right). Statistical comparisons were determined using ANOVA and post-hoc test with Tukey correction (* p < 0.05, ** p < 0.01, *** p < 0.001, and **** p < 0.0001).

Comparing the relative cell occurrences at P20 showed on average an enrichment of OPC (72 ± 2%), COP (33 ± 5%), NFOL1 (59 ± 3%), MOL1 (76 ± 4%), MOL5 (36 ± 4%) and MOL6 (48 ± 9%) in GM compared to WM in the spinal cord. In contrast, MOL2 (54 ± 7%) and MOL3 (79 ± 7%) were enriched in WM (p = 0.0003, p = 0.01, p = 0.002, p = 0.002, p = 0.003, p = 0.02, p = 0.02, p = 0.0009, respectively; n=6 tissue sections per condition, **Figure 5C, left**). These distributions were not substantially altered in P60 spinal cord, with an average enrichment of OPCs (60 ± 7%), COP (29 ± 7%), MOL1 (70 ± 6%) and MOL6 (59 ± 8%) in GM compared to WM and reversibly an average enrichment of MOL2 (80 ± 6%) and MOL3 (92 ± 1%) in WM compared to GM (p = 0.002, p = 0.04, p = 0.003, p = 0.007, p = 0.005, p = 0.002, respectively; n=6 tissue sections per condition, **Figure 5C, left**). MOL2/3 and MOL5/6 are also densely present in the dorsal funiculi of WM (**Figure 5B**). Thus, in contrast to the brain, we did not observe major differences in the pace of differentiation/myelination between P20 and P60. Nevertheless, there is a clear spatial preference for MOL2 in WM and MOL5/6 in GM at P20 and P60, as we have previously reported (Floriddia et al, 2020).

While there is a decrease with age in the GM of OPC (39 ± 2%), COP (69 ± 3%), MFOL1 (70 ± 3%), MOL2 (60 ± 6%), there is an increase of MOL5 (23 ± 3%) and MOL6 (54 ± 1%) (p = 0.002, p = 0.001, p = 0.03, p = 0.008, p = 0.008, p = 0.00013, respectively; n=6 tissue sections per condition, **Figure 5C, right**). In the WM, a similar pattern was observed for MOL6 (37 ± 7% increase). However, in contrast to GM, OPC, COP, MFOL1, MOL2 and MOL5 were not significantly altered in WM (p = 0.05; n=6 tissue sections per condition, **Figure 5C, right**). Also, in contrast to the brain, MOL1 was not reduced from P20 to P60 in the spinal cord. Thus, there is a region specificity in the dynamics and abundance of OL populations in the WM and GM of different regions of the CNS.

Comparing P20 GM directly for brain vs. spinal cord, showed increases for NFOL1 (42 ± 7%), NFOL2 (53 ± 13%), MFOL1 (55 ± 8%) and MFOL2 (58 ± 9%) in the brain and increases of MOL1 (24 ± 4%), MOL2 (33 ± 9%), MOL5 (54 ± 8%) and MOL6 (91 ± 1%) in the spinal cord (p = 0.02, p = 0.05, p = 0.04, p = 0.05, p = 0.01, p = 0.05, p = 0.03, p = 0.00003, respectively; n=6 tissue sections per condition, **Figure 6A, left**), indicating that the GM in the spinal cord is in a more advanced maturation state than in the brain. A similar pattern is found in the WM, where at P20 there is an increase of OPC (60 ± 7%), COP (48 ± 8%), NFOL1 (79 ± 3%) and MOL1 (64 ± 6%) in the brain and an increase of MOL2 (78 ± 1%), MOL3 (54 ± 9%), MOL5 (55 ± 1%) and MOL6 (90 ± 2%) in the spinal cord (p = 0.04, p = 0.05, p = 0.002, p = 0.01, p = 0.0005, p = 0.02, p = 0.006, p = 0.01, respectively; n=6 tissue sections per condition, **Figure 6A, right**).

**Figure 6:**
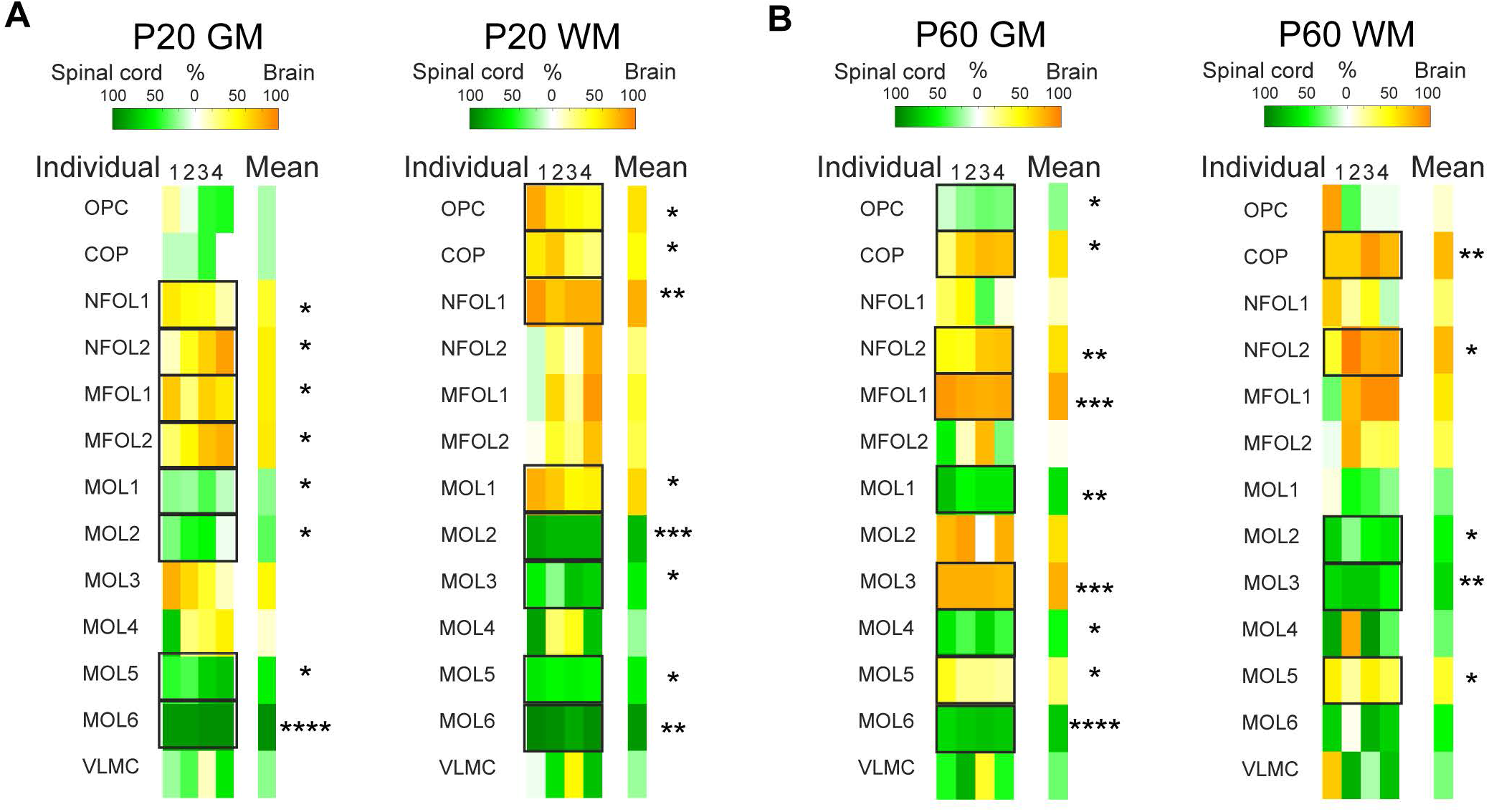
Change of relative oligodendrocyte abundances in P20 and P60 GM and WM for brain vs. spinal cord **a**, Change of relative abundances of oligodendrocytes between brain and spinal cord for P20 GM (left) and P20 WM (right). Statistical comparisons were determined using ANOVA and post-hoc test with Tukey correction (* p < 0.05, ** p < 0.01, *** p < 0.001, and **** p < 0.0001). **b**, Change of relative abundances of oligodendrocytes between brain and spinal cord for P60 GM (left) and P20 WM (right).

At P60, GM showed increases of COP (61 ± 10%), NFOL2 (60 ± 6%), MFOL1 (83 ± 2%) and MOL3 (79 ± 1%) in the brain and increases of OPC (22 ± 3%), MOL1 (60 ± 4%), MOL4 (49 ± 5%) and MOL6 (70 ± 2%) in the spinal cord (p = 0.03, p = 0.006, p = 0.0003, p = 0.0003, p = 0.02, p = 0.001, p = 0.03, p = 0.00001, respectively; n=6 tissue sections per condition, **Figure 6B, left**). P60 WM shows increases of COP (75 ± 4%), NFOL2 (75 ± 10%) and MOL5 (41 ± 7%) in the brain and increases of MOL2 (50 ± 8%) and MOL3 (63 ± 4%) in the spinal cord (p = 0.004, p = 0.02, p = 0.02, p = 0.03, p = 0.007, respectively; n=6 tissue sections per condition, **Figure 6B, right**). Interestingly, at P60, MOL2 is enriched in the WM spinal cord versus the WM brain, but the opposite seems to apply to the GM. MOL1 and MOL6 appear more enriched in the spinal cord, while MOL5 in the brain at P60. Thus, at P60, region specific differences on the abundance of the mature OL populations become clearer and consolidated.

### Oligodendroglia populations’ neighboring preferences are altered from the juvenile to the adult CNS

As OL populations show different spatial preferences in brain and spinal cord, the question arises if certain OL populations are organized in distinct neighborhoods. Therefore, we performed a nearest neighbor analysis, where each cell is represented as a node in a graph. For each node there is a corresponding region consisting of all points of the plane closer to that node than to any other. This mathematical approach partitions the plane into regions and is called Voronoi tessellation (see shades in **Figure 7A and D**). Each node is then connected to all its neighboring nodes (Delaunay triangulation) and the distance between the points can be measured using the Euclidean distance. The neighbor with the closest Euclidean distance is the nearest neighbor. We calculated the nearest neighbor distances for each OL population for P20 GM and WM as well as P60 GM and WM and visualized the distribution of relative abundances of the nearest neighbors in heatmaps. Data are normalized to account for the different absolute abundances of the different OL populations.

**Figure 7:**
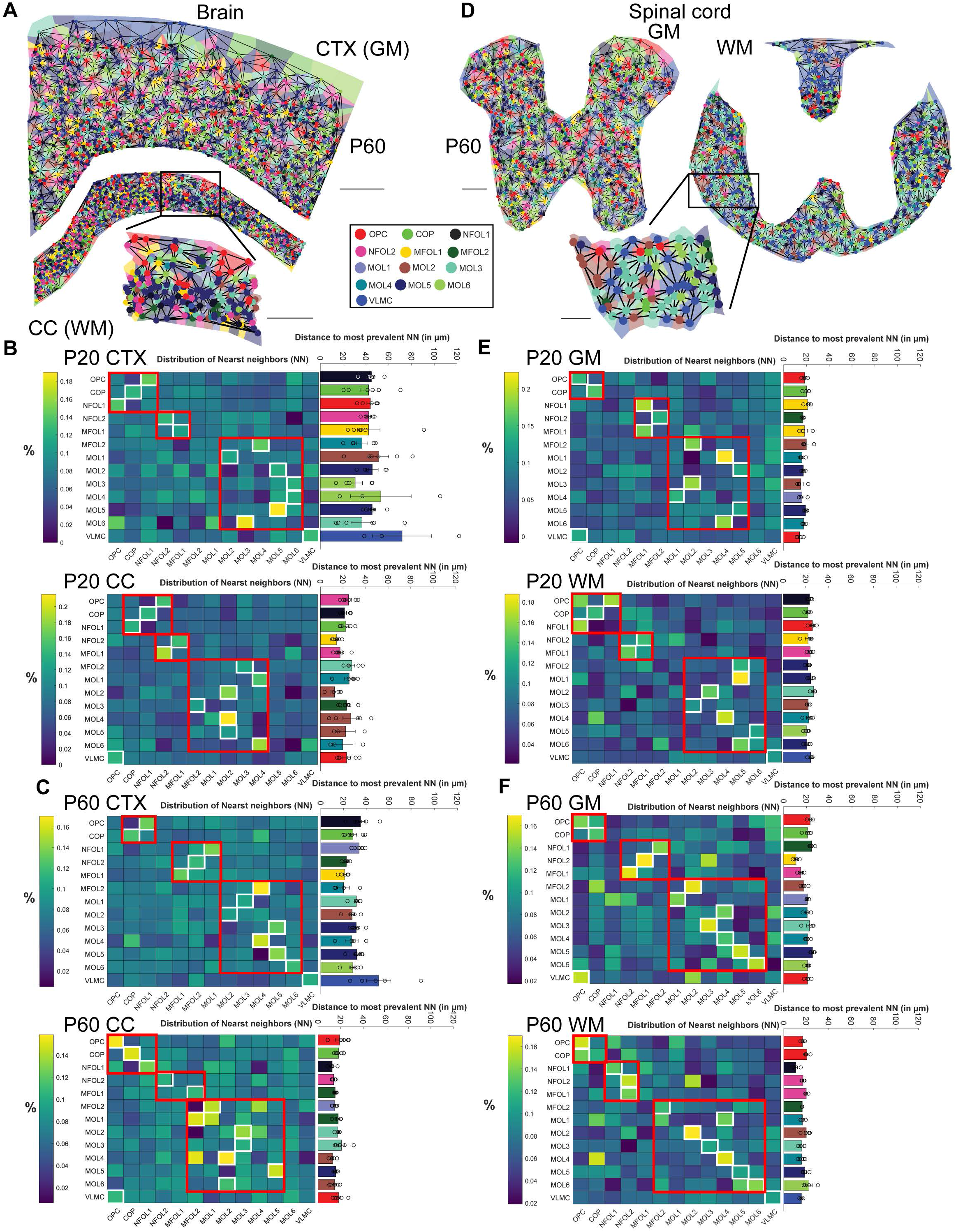
Nearest neighbor (NN) analysis for oligodendrocytes **a**, For each cell a region is plotted (tiles in the background), where all points of the region are closer to that cell than to any other (Voronoi tessellation). Each cell is a node in the graph and the edges connect the neighboring cells (Delaunay triangulation). A zoom-in is shown for the WM region. The main scale bar is 500 μm, the zoom-in scale bar is 250 μm. **b**, Heatmaps of the nearest neighbor likelihood (most probable NN per oligodendrocyte type highlighted by white square) and barplot of average distances to most prevalent NN for P20 CTX (top) and CC (bottom). **c**, Same as **b** for P60 CTX and CC. **d**, Same as **a** for spinal cord. The main scale bar is 100 μm, the zoom-in scale bar is 50 μm. **e**, Same as **b** for spinal cord P20 GM and WM. **f**, Same as **e** for spinal cord P60 GM and WM.

We mainly found three patterns of neighborhood, corresponding to different stages during OL lineage progression, and that were largely consistent over the different conditions (**Figure 7B-F**). Pattern I corresponded to early OL differentiation. In the P20 brain, we observed that OPCs preferentially neighbor with NFOL1 (P20 CTX) or NFOL2 (P20 CC), COP with COP (P20 CTX) or NFOL1 (P20 CC). Pattern II included NFOL2 neighboring with NFOL2 (P20 CTX) or MFOL1 (P20 CC) and MFOL1 with MFOL1 (P20 CTX) or NFOL2 (P20 CC), corresponding to a transition to a myelinating stage (**Figure 7B, top**). Pattern III includes the last stages of myelination and neighborhoods between mature OLs. MFOL2 was found to have MOL4 (P20 CTX) and MOL3 (P20 CC) as its preferential nearest neighbors and mature populations tend to group with one another, although with dissimilarities between CC and CTX (**Figure 7B, bottom**). The average Euclidean distances between OL populations was not significantly different in the P20 CTX, being between 40-50 μm. However, the distances are more variable in the CC, ranging from 15 μm MOL2-MOL2 to nearly 30 μm for MFOL2-MOL3.

P60 CTX has similar patterns, although with different combinations in pattern II, with NFOL2 neighboring MFOL2, while NFOL1 neighboring MOL1 (**Figure 7B, top**). Interestingly, we found OLs often neighboring OLs of the same type, such as COP, MFOL1, MOL2, MOL4, MOL5 and MOL6 in P60 CTX. This pattern is also present to some extent in P60 CC, for MOL1, MOL3 and MOL5, but was even more striking with OPC, COP, NFOL1, NFOL2, (**Figure 7C, bottom**). This homotypic neighborhood structure in adulthood might reflect a stabilization of the OL phenotypes, due to a reduction of migration of OPCs and stable myelin wrapping of defined neuronal populations in the adult brain. This homotypic neighborhood is already observed to a higher degree in the spinal cord at P20 (**Figure 7E**), being further stabilized at P60 (**Figure 7F**). The distances between OL populations ranged from 10-25 μm, similar to brain WM, but not brain GM.

VLMCs show preference to other VLMCs or to OPCs in the P20 CC, P60 CC, P20 GM and P60 GM **(Figure 7B-F)**. VLMCs are associated in the vasculature and can thereby be a proxy for vasculature proximity at the CNS parenchyma. OPCs have been reported to be closely associated with the vasculature during embryonic development, during their migration (Tsai et al, 2016). While OPC migration in the adult CNS is limited, it is reactivated upon injury, alongside OPC association with the vasculature (Niu et al, 2019). Thus, our nearest neighbor analysis supports the association of OPCs with the vasculature.

## DISCUSSION

In this study, we analyzed simultaneously the expression of 124 genes expressed in different OL lineage populations in three regions of the juvenile and adult CNS. *In situ* sequencing (ISS) analysis allowed us to determine the spatial distribution of OLs with an unprecedented resolution.

The distribution of OL lineage cells in the forebrain WM and GM at P20 and P60 suggests that differentiation and myelination advances earlier in CTX (GM) than in CC (WM) in the brain. We observed at P20 an enrichment of COPs in WM, while more intermediate stages as NFOL2 and MFOL1/2 were enriched in the GM (**Figure 3A**). This data suggests that differentiation is actively ongoing at this stage in the CC, while in CTX OL are rather in a myelination phase. Indeed, using lineage tracing with the NG2creERTMBAC:ZEG mice, Zhu and colleagues showed that OL differentiation plateaus between P10 and P60 in the CTX, while it is still occurring until P60 in the CC (Zhu et al, 2011). Interestingly, these intermediate myelin-forming states are enriched in the CC WM at P60, while the CTX GM has already transitioned to fully mature OLs (**Figure 3A**). This is consistent with previous lineage tracing findings with various transgenic mouse lines, indicating that OPCs in the cortical GM have a longer cell cycle time than in WM CC at P20 and P60 (Young et al, 2013; Psachoulia et al, 2009), and that differentiation and myelination of the CC WM can extend until at least 240 days in the mouse, but to a lesser extent in the CTX GM (Sturrock, 1980; Rivers et al, 2008; Kang et al, 2010; Zhu et al, 2011).

The composition of mature OL populations in the P60 WM and GM in the brain is quite similar, with a marked increase in MOL6 and MOL5, which becomes the prevalent population in these regions (**Figure 2 and Figure 3**). In contrast, the spinal cord presents clear spatial preferences of oligodendroglial populations, which are maintained from P20 to P60 (enrichment of OPC/COP, MOL1 and 5/6 in GM and MOL2/3 in WM). It will be interesting to investigate in the future whether such spatial preference is established later on in adulthood in the brain. Alternatively, it is possible that other regions in the brain (for instance in ventral regions) exhibit spatial preferences.

We observed a decrease in MOL1 consistent with our previous RNASCOPE dataset, where there was a tendency for decrease in the MOL1 population with age (Floriddia et al, 2020), which becomes clear with ISS. MOL1 is a population we previously observed to be dramatically increased at the injury site upon spinal cord injury (Floriddia et al, 2020). The observed developmental decrease in MOL1 occurs as populations such as NFOLs and MFOLs are also reduced, not only as a result of the dynamics of differentiation between the two time points, but also due to programmed cell death (Raff et al, 1993; Barres et al, 1992; Trapp et al, 1997; Sun et al, 2018). The presence of MOL1 in niches during development and pathology that are associated with cell death could suggest that this mature OL state might be associated with stress and even apoptosis. Indeed, MOL1 is characterized by the expression of genes such as *Egr1*, *Egr2*, *Fos*, *Arc*, among others (Marques et al, 2016). Many of these genes can be upregulated upon neuronal activation, but also upon stress and apoptosis (Hrvatin et al, 2018). However, stress and apoptosis can be induced by the processing of tissue in single cell transcriptomic approaches (Brink et al, 2017; Marsh et al, 2020). Thus, it is challenging to determine whether stress/activated related signatures are truly biological or artefactual. Nevertheless, the detection of MOL1 *in situ* in the CNS by technologies such as RNASCOPE (Floriddia et al, 2020) and ISS (this study) strongly suggests that this population represents a physiological state of mature oligodendrocytes, although it remains unclear if it is a stress-related transient state, which might become stabilized in certain developmental or pathological niches.

One of the major advantages of technologies such as ISS is the spatial resolution, and the possibility to examine population distributions in anatomical areas. We superimposed our data with previously defined cortical areas. Interestingly, we observed newly-formed and myelin-forming oligodendrocytes present mainly at the layers V-VII of the cingulate 1 and motor 1 cortical areas at P20, while MOL5 was already specified specifically at the somatosensory cortex. In contrast, there was a rather uniform distribution of the OL lineage populations at P60. Thus, our data indicates a medial-to-lateral gradient of OL lineage progression at P20 in the CTX, and possibly a deep-to-superficial layer gradient, that is resolved at P60. While most of the mature OL lineages has been previously shown to accumulate in the deeper layers of the cortex (McGee et al, 2005; Vincze et al, 2008; Hughes et al, 2018), the medial-to-lateral gradient was unexpected, and suggests that OL differentiation and myelination occur at different time points at different areas in the cortex. Such arealization of myelination has been shown at the mouse corpus callosum between P3 and P21, with sensory pathways being myelinated first, followed by motor pathways and association pathways (Vincze et al, 2008), and also in humans (Kinney et al, 1988; Brody et al, 1987). Interestingly, specific mature OL populations have been shown to myelinate inhibitory neurons, excitatory neurons, or both (Zonouzi et al, 2019), although it is unclear if these populations have any correspondence to the transcriptional states as MOL1-6. It is possible that neurons and other cells such as microglia present in layers V-VII might have a particular role on the differentiation, myelination and final subtype specification of MOLs. Neuropilin-1 has been recently shown to regulate OPC proliferation and to be expressed only in activated microglia in the developing WM (CC), but not in GM (CTX) or adult CC (Sherafat et al, 2021). It is also possible that distal neuronal connections modulate the timing of OL lineage progression in the cortex, as many of these cortical areas project to distinct and remote areas of the CNS.

We also observed that the maturation into MOLs is more advanced at P20 in both WM and GM of the spinal cord, when compared to the brain (**Figure 6**). This is consistent with previous reports of an posterior-to-anterior gradient of myelin glycoprotein (Mog) expression in the post-natal rat CNS (Coffey & Mcdermott, 1997), autopsy studies in humans (Brody et al, 1987), longer cell cycle time of OPCs in the P21 and P60 spinal cord WM compared to corpus callosum (Young et al, 2013) and reduced capacity of differentiation of OPCs at the P30 in the spinal cord when compared to the brain (Sturrock, 1980; Rivers et al, 2008; Kang et al, 2010; Zhu et al, 2011).

In our previous study, based on RNASCOPE, MOL2 cells could barely be detected in CTX and CC and were prominent in the spinal cord (Floriddia et al, 2020). Using several markers for MOL2, we could confirm the prevalence of this population in the spinal cord WM versus brain WM (**Figure 6B**), but could now detect a higher number of this population in the brain, and an increase in these brain regions from P20 to P60 (**Figure 3B**). This suggests that MOL2 might also have relevant functions not only in the spinal cord, but also in specific areas of the brain, which could be particularly relevant given that we observed that MOL2 is particularly affected in the context of injury (Floriddia et al, 2020).

Another important aspect to consider is that our neighborhood analysis is centered on the nucleus of each cell. OPCs and OLs exhibit extensive processes, which sense the environment and neighboring cells. Thus, while the cell body of certain cells might be distant, it is possible that their processes are in close contact. Interestingly, our nearest neighbor analysis indicates that the closest neighbors of OPCs are VLMCs, or OPCs themselves (**Figure 7**). However, OPCs have been shown to self-avoid in the mouse adult (P60-P90) somatosensory CTX, thereby maintaining distinct territories (Hughes et al, 2013). This apparent paradox most likely reflects the high level of area coverage of OPCs in the CNS, as observed in Figures 4 and 5. Cell death and differentiation of OPCs is compensated by proliferation and migration of neighboring OPCs, maintaining thus a constant and high density (Hughes et al, 2013; Buchanan et al, 2021) and remodeling (Xiao et al, 2021), and response to inflammatory insults in the context of injury or disease (Falcão et al, 2018).

Our ISS dataset provides the first detailed analysis of oligodendrocyte lineage spatial organization in the mouse juvenile and adult WM and GM. While confirming previous literature findings regarding the caudal-rostral timing of myelination and association of OPCs with the vasculature, our data extends these findings by giving a detailed view of the spatial organization of distinct OL lineage populations. Furthermore, it uncovers the timing of OL differentiation and myelination in specific areas of the cortical GM, and OL lineage neighborhoods. Future integration of our dataset with other emerging spatial transcriptomics targeting in detail subtypes of neurons, astrocytes, microglia, among others, is bound to lead to a better understanding of the cellular architecture of the CNS and the principles by which it is established.

## MATERIALS AND METHODS

### Animals

All animals (Pdgfrα::CreERTM-loxP-RCE (Z/EG) (Kang et al, 2010) with mixed C57BL/6NJ and CD1 background) were free from the most common mouse viral pathogens, ectoparasites, endoparasites and mouse bacterial pathogens harbored in research animals. The battery of screened infective agents met the standard health profile established in Karolinska Institutet animal housing facilities. Mice were kept with the following light/dark cycle: dawn 6:00–7:00, daylight 7:00–18:00, dusk 18:00–19:00 and night 19:00–6:00 and housed to a maximum number of five per cage in individually ventilated cages (IVC Sealsafe GM500, Tecniplast). Cages contained hardwood bedding (TAPVEI), nesting material, shredded paper, gnawing sticks and a cardboard box shelter (Scanbur). The mice received a regular chew diet (either R70 diet or R34, Lantmännen Lantbruk, or CRM-P, 801722, Special Diet Services). General housing parameters such as relative humidity, temperature and ventilation follow the European convention for the protection of vertebrate animals used for experimental and other scientific purposes treaty ETS 123. Briefly, consistent relative air humidity of 50%, 22 °C and the air quality is controlled with the use of stand-alone air handling units supplemented with high efficiency particulate air-filtered air. Husbandry parameters were monitored using ScanClime (Scanbur) units. Water was provided in a water bottle, which was changed weekly. Cages were changed every other week. All cage changes were done in a laminar airflow cabinet. Facility personnel wore dedicated scrubs, socks and shoes. Respiratory masks were used when working outside of the laminar airflow cabinet. All experimental procedures in this study were conducted in accordance with the European directive 2010/63/EU, local Swedish directive L150/SJVFS/2019:9, Saknr L150 and Karolinska Institutet complementary guidelines for procurement and use of laboratory animals, Dnr 1937/03-640. The procedures described here were approved by Stockholms Norra Djurförsöksetiska nämnd, the local committee for ethical experiments on laboratory animals in Sweden, lic.nr. 130/15, 144/16, and 1995/2019.

### *In situ* sequencing assay

P20 and P60 mouse brain tissues and spinal cord tissues were snap frozen in OCT mounting medium, coronally sectioned in 10 μm-thick cryosections and stored at −80°C until fixation. 6 coronal brain sections (Bregma 1.10 mm, 3x P20, 3x P60, hemisected) and 8 thoracic spinal cord sections (4x P20, 4x P60) were studied. The brain sections were hemisected. *In situ* sequencing, a targeted multiplexed mRNA detection assay employing padlock probes, rolling circle amplification (RCA) and barcode sequencing, was applied. Padlock probes targeting a set of 124 genes and the CartaNA NeuroKit (1010-01, CartaNA AB, now part of 10x Genomics) were used (similar to (Soldatov et al, 2019)). For a step-by-step protocol of the *In situ* sequencing chemistry see (Hilscher et al, 2020). In short, 10 μm-thick cryosections were fixed by 3% (w/v) paraformaldehyde (PFA) in DEPC-treated PBS (DEPCPBS) at room temperature (RT) for 5 min, and they were washed in DEPC-PBS followed by 0.1N HCl treatment at RT for 5 min. The sections were washed in DEPC-PBS again, and they were subjected to reverse transcription, probe ligation and rolling circle amplification reactions. The RCA products were detected by hybridization of AF750 labelled detection oligo, and decoded through four cycles of barcode sequencing, each involving a base-specific incorporation of fluorescence dyes (A=Cy5, C=Texas Red, G=Cy3, T=AF488), imaging, and removing the incorporated dyes.

### Imaging setup and image analysis

Images were acquired using a Zeiss Axio Imager Z2 epifluorescence microscope (Zeiss Oberkochen, Germany), equipped with a 40x objective. A series of images (10% overlap between two neighboring images) at different focal depths was obtained and the stacks of images were merged to a single image thereafter using the maximum-intensity projection (MIP) in the Zeiss ZEN software. After exporting images in .tif format, images were aligned between cycles and then stitched together using the MIST algorithm. The stitched images were then re-tiled into multiple smaller images, henceforth referred to as tiles. Tiled DAPI images were segmented with standard watershed segmentation. The tiled images with RCA products were top-hat filtered, followed by initial nonlinear image registration, spot detection, fine image registration, crosstalk compensation, gene calling and cell calling using the pciSeq pipeline (https://github.com/acycliq/full_coronal_section, (Qian et al, 2019)).

### Genes and cell typing

The analysis pipeline further assigns genes to cells and then cells to cell types. It leverages the area, the global x and y coordinates and the unique cellular identifier of the segmented cells as well as the global x and y coordinates of the genes. The assignment is done using a probabilistic framework based on single-cell RNA sequencing data. Here, our targeted genes were selected from single-cell RNA sequencing data from (Marques et al, 2016) based on median expression values to target major markers for the oligodendrocyte populations. Cell annotations and definitions used in pciSeq are also from (Marques et al, 2016). Following these definitions pciSeq then represents the cells as pie charts, with the angle of each slice proportional to the cell type probability.

## Data analysis

The highest probability in the pie charts was used to define the final oligodendrocyte type (winner-takes-all principle). Only cells containing at least one Plp1 gene were kept for downstream analysis. The downstream analysis was carried out with custom-made scripts in Matlab and Python (https://github.com/Moldia). Comparisons of cell counts between conditions (CTX and CC or P20 and P60) were calculated by normalizing the cell counts for each sample first. Thereby, the cell counts for each oligodendrocyte population were divided by the total number of Plp1+ cells per sample in order to account for differences in sample size and to obtain relative cell counts (values between 0 and 100). The ratios of the relative cell counts were then calculated as *ratio = 100/(condition X for cell_type A/condition Y for cell_type A)* with *condition X* denoting the larger cell count and *condition Y* denoting the smaller cell count (*X, Y = {CTX, CC, P20, P60}*). The ratio was subtracted from 100 to express increases, i.e. if *condition X* is 30 and *condition Y* is 10 the heatmap shows a value of 66.66%, meaning that the increase from *condition Y* to *condition X* is 20 (or 66.66% of *condition X*). Bar plots show mean ± SEM (standard error of mean). Density plots were generated using the Matlab function ksdensity and registered to Allen Mouse Brain Atlas using a custom-made script similar to (Partel et al, 2020). Statistical comparisons were determined using two-tailed Student’s paired t test, and to account for multiple comparisons, the data were analyzed using ANOVA and post-hoc test with Tukey correction (* p < 0.05, ** p < 0.01, *** p < 0.001, and **** p < 0.0001).

## Acknowledgements

We thank Elisa Floriddia for the preparation of tissue sections and the help with the marker selection. We acknowledge Amitha Raman and the In situ sequencing facility unit, part of the spatial and single cell biology platform at Science for Life Laboratory, Sweden for technical assistance with the *in situ* sequencing experiments. We thank Mats Nilsson lab members for their insights and comments. We would like to thank T. Jimenez-Beristain for writing the laboratory animal ethics permit 1995_2019 and assistance with animal experiments, and the staff at Comparative Medicine-Biomedicum.

## Author contributions

M.M.H. designed and performed the experiments, analyzed and interpreted the data and wrote the manuscript. C.M.L. and P.K. analyzed and interpreted the data. C.Y. performed experiments. M.N. and G.C.B. wrote the manuscript. All authors read the manuscript and provided feedback.

---

**Figure S1:**
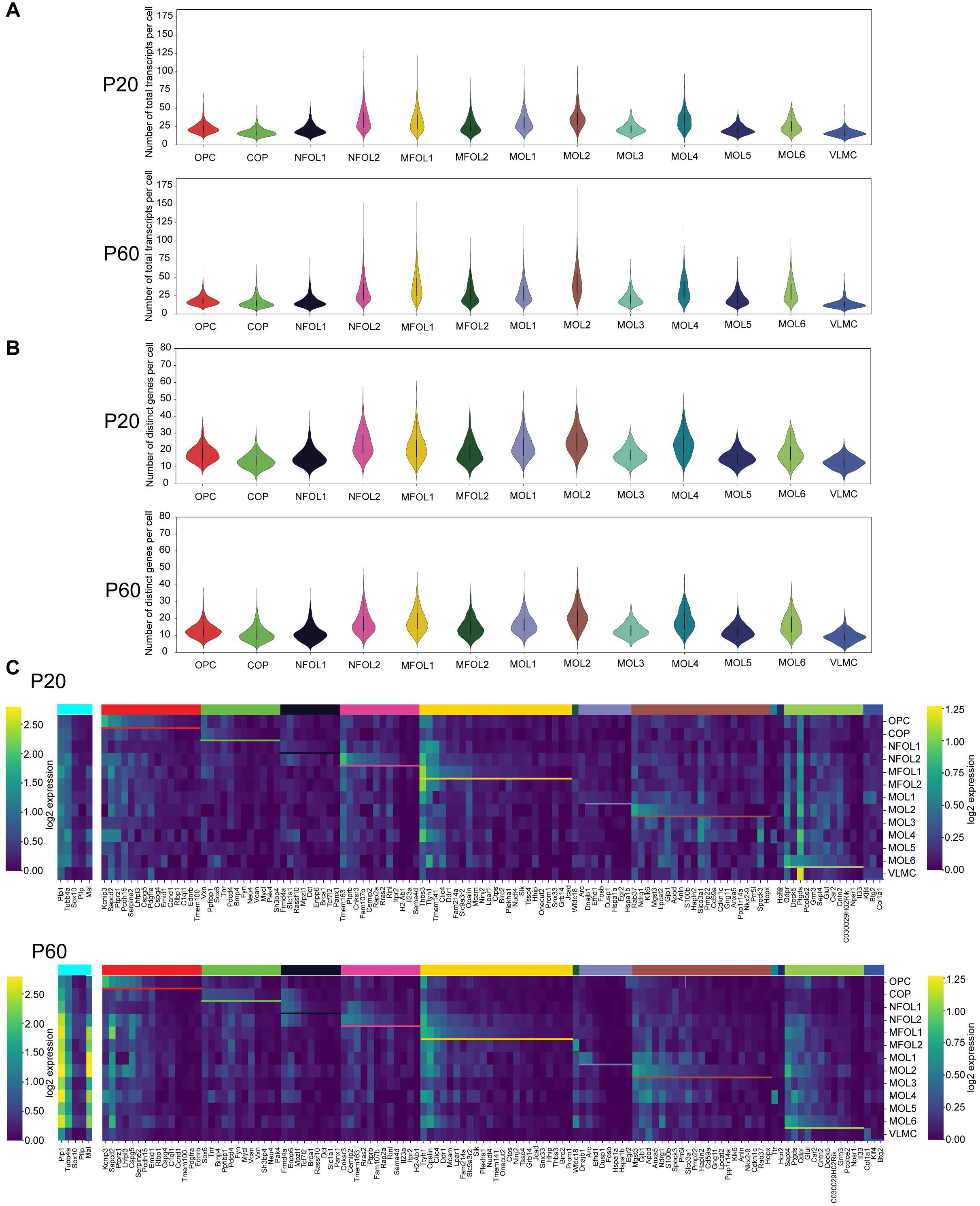
Genes per cell for P20 and P60 oligodendrocytes **a**, Violin plots of the number of total transcripts detected in the oligodendrocyte lineage cells and VLMCs with targeted *In Situ* sequencing (top: P20; bottom: P60; CTX and CC joined). **b**, The number of distinct genes in oligodendrocytes (top: P20; bottom: P60; CTX and CC joined). **c**, Mean log2-transformed expression of the targeted genes in oligodendrocyte lineage cells and VLMCs, grouped by cell type (see bar on top) for P20 and P60, respectively. The cyan bar denotes pan markers for oligodendrocytes. Note that the scale in the left and right color bars are different.

**Figure S2:**
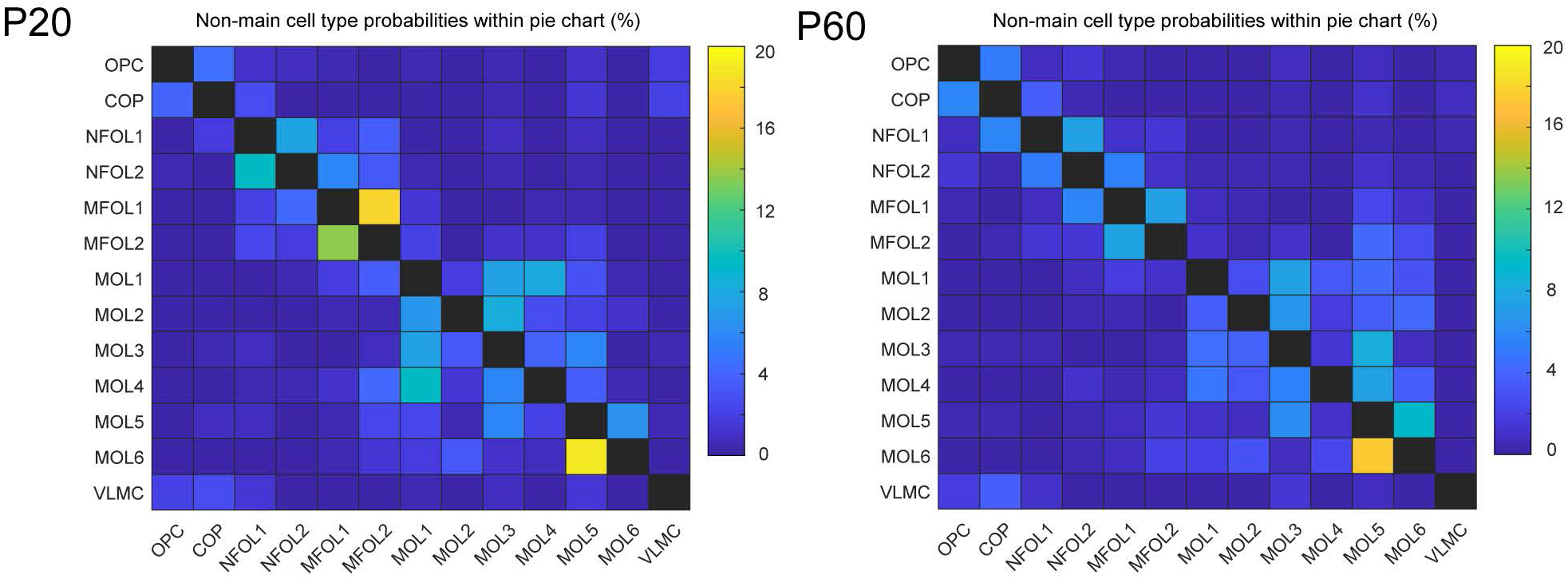
Probabilities of cell types for P20 and P60 Heatmap of the probability distribution in the pie charts that are not the highest probability (mean confusion matrix; left: P20; right: P60; CTX and CC joined).

**Figure S3:**
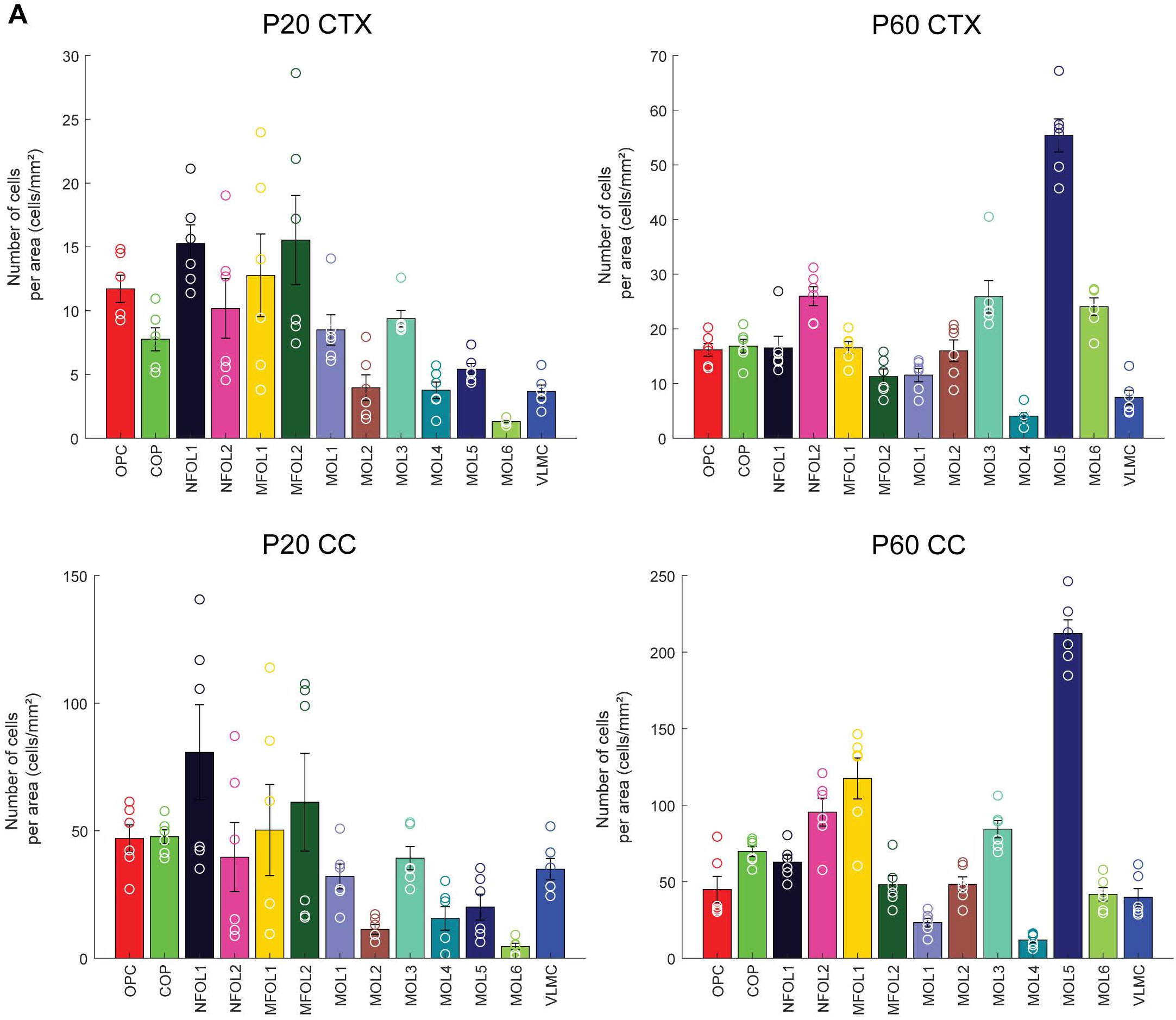
Oligodendrocyte density per area for P20 and P60 CTX (GM) and CC (WM) **a**, Top: Number of oligodendrocytes per area (in cells/mm^2^) for CTX (top) and CC (bottom). **b**, Same as **a** for P60.

